# Infants are superior in implicit crossmodal learning and use other learning mechanisms than adults

**DOI:** 10.1101/139535

**Authors:** Sophie Rohlf, Boukje Habets, Marco von Frieling, Brigitte Röder

## Abstract

While adults have to continuously adapt their internal representations of the sensory world, infants need to first acquire these models. We used event-related potentials to test the hypothesis that infants extract crossmodal statistics implicitly while adults learn them when task relevant. Six-month-old infants and adults were passively exposed to frequent standard audio-visual combinations (A1V1, A2V2, p=0.35 each), rare recombinations of the standard stimuli (A1V2, A2V1, p=0.10 each), and a rare deviant audio-visual combination with an infrequent auditory and visual element (A3V3, p=0.10). While both infants and adults differentiated between rare deviants and standards at early processing stages, only infants discriminated standards from recombined stimuli at a later processing stage. A second experiment revealed that adults discriminated recombined from standard combinations only when crossmodal combinations were task relevant. These results demonstrate a heightened sensitivity for crossmodal statistics in infants and a change in learning mode from infancy to adulthood.

## Introduction

After birth infants are immediately exposed to a sensory world comprising input of multiple sensory modalities. The developing brain must adapt to the statistical properties of the sensory environment (Fiser et al., 2010) since genetically defined neural circuits are usually crude. Indeed a high sensitivity of infants to statistical regularities within single sensory systems has often been demonstrated (Fantz, 1964; Saffran et al., 1996; Fiser & Aslin, 2002; Bulf et al., 2011). The seminal study of Saffran et al. (1996) reported that eight-month-old infants quickly learn transitional probabilities between syllables by pure exposure to an artificial language. This ability was interpreted as a basic mechanism allowing infants to segment a language. Similar results were found for non-linguistic auditory sequences and for visual patterns (Fiser & Aslin, 2002), demonstrating a modality independent sensitivity of infants to statistical patterns in their sensory environment which moreover is not unique to linguistic material. For example, in the visual domain, there is strong evidence that infants are able to implicitly learn subtle statistical relationships among visual objects (Fiser & Aslin, 2002; Bulf et al., 2011; Kirkham et al., 2002). Nine-month-old infants who were exposed to multi element visual scenes, showed greater interest in element pairs which co-occurred more frequently than in pairs which co-occurred less frequently. Moreover, the infants were sensitive to the predictability between elements of the pairs as manifested by the conditional probability relations between these elements (Fiser & Aslin, 2002). Infants' ability to extract statistical patterns of visual stimuli was found even in younger age groups (Kirkham et al., 2002); two-, five-, and eight-month-old infants were habituated to sequences of discrete visual stimuli whose ordering followed a statistical predictable pattern. Subsequently the infants were shown the previously encountered pattern alternating with a novel pattern of identical stimulus components. All age groups looked longer at the novel sequences providing evidence for the detection of visual statistical regularities at an early developmental stage. These results suggest that infants own powerful mechanisms for extracting the statistical properties of their sensory input without any instructions, explicit feedback, or intentional awareness (Lany & Saffran, 2013; Krogh et al., 2013).

The ability of infants to detect crossmodal statistical regularities within their sensory environment is less well understood, but some basic multisensory abilities, such as multisensory temporal synchrony detection seem to exist within the first month of life (Lewkowicz, 1992). In the next months the capability to perceive higher-level and more complex multisensory relations starts to develop. For example, at the age of six months infants were shown to perceive duration-based (Lewkowicz, 1992) and spatio-temporal based crossmodal relations (Scheier et al., 2001). Furthermore, there is evidence that similar to adults, infants take advantage of crossmodal events in terms of a better discrimination and a faster responsiveness to bimodal compared to unimodal information (Bahrick et al., 2004; Lewkowicz & Kraebel, 2004). First evidence for multisensory facilitation was found in eight-month-old infants as indicated by faster eye movements to spatially aligned auditory and visual cues compared to eye movements to each of these stimuli alone (Neal et al., 2006). Moreover, other studies revealed multisensory benefits for perceptual learning in infants (Bahrick & Lickliter, 2002; Frank et al., 2009). Five-month-old infants were habituated to either an audio-visual rhythm or the same rhythm presented unimodally. In the crossmodal condition, infants were able to discriminate between the familiar and a novel rhythm, whereas no discrimination was observed for the unimodal stimuli (Bahrick & Lickliter, 2002). Corresponding results were found for the learning of an abstract rule in five-month-old infants: they were able to learn the sequence if defined by redundant visual shapes and speech sounds but not if only one sensory modality was involved (Frank et al., 2009). These results suggest that infants are able to learn and use associations between auditory and visual stimuli. However, it must be taken into account that the multisensory effects in infants were not tested against statistical facilitation (probability summation, see Miller, 1982).

Several studies on crossmodal association learning have reported that infants at the age of three months, but not younger, are able to learn specific voice-face pairings; infants were habituated to different unfamiliar voice-face pairings. In the post-familiarization test the infants showed higher attention to the learned voice-face pairings as compared to the novel combinations. The latter category comprised a voice and a face they had heard and seen previously, but the combination of the voice and face was new (Brookes et al., 2001; Bahrick et al., 2005). More recently, near-infrared spectroscopy (NIRS) and event-related potentials (ERPs) were used to test whether infants are able to learn crossmodal associations between arbitrary auditory and visual stimuli. Emberson et al. (2011) used an audio-visual omission paradigm with six-month-old infants and found similar visual cortex activation for an auditory stimulus as well as visual stimuli that had been previously combined with this auditory stimulus. The authors interpreted their findings as evidence for top-down mechanisms to be in place as early as six month of age. Kouider et al. (2015) exposed twelve-month-old infants to pictures of faces paired with one sound and pictures of flowers paired with another sound. During the test phase the sound preceded the visual stimulus and was either congruent or incongruent with the learned combinations (additionally no sound was used in one third of the trials). An enhanced early negative ERP for congruent visual stimuli as well as an enhanced late positive ERP for incongruent visual stimuli were found. Both studies demonstrate that infants are able to learn crossmodal combinations to which they were exposed. However, none of these studies used an adult control group. Thus, it remains an open question whether developmental and adult crossmodal learning recruit the same mechanisms. In this context it is interesting to notice that Janacsek et al. (2012) demonstrated superior implicit statistical learning of visual sequences in young children compared to older children and adults; a follow-up study indicated that this advantage was lost when they became more reliant on explicit learning (2013).

Based on animal studies it has been proposed (Keuroghlian & Knudsen, 2007) that developmental and adult plasticity, and thus learning, differ due to different brain states particularly during the sensitive phase molecular mechanisms dominate that allow for quick and extensive functional and structural synaptic plasticity (synaptogenesis, synaptic strengthening and elimination) as well as for the emergence of the functional adaptive connectivity. By contrast, in adulthood these functionally tuned and to some degree stabilized neural circuits undergo adaptations when relevant to the system. These age dependent changes from developmental to adult plasticity are impressively demonstrated by a study on auditory cortex plasticity in rats: while passive exposure to sounds of a specific frequency results in a permanent reorganization of auditory cortex during the sensitive phase, adult rats reorganize only those aspects of the auditory cortex that are task relevant: for example, rats were exposed to sounds which varied both in sound frequency and level. When they had to discriminate them with respect to sound frequency the frequency representation of auditory cortex changed while the level representation changed when level rather than sound frequency was task relevant (de Villers-Sidani et al., 2007). These findings suggest that adult learning seems to depend to a larger degree on attention and context such as task relevance and reward expectations (Keuroghlian & Knudsen, 2007; Bavelier et al., 2010). This hypothesis was supported by Riedel and Burton (2006) who investigated whether learning of auditory sequences is influenced by task demands; when using a serial reaction time task related to the feature of the auditory stimulus, they found learning effects in adult participants while a passive exposure did not result in learning. Similarly, the statistical relations of concurrently presented visual streams were only learned by adults for the attended and not for the unattended streams (Turk-Browne et al., 2005).

In the present study we investigated multisensory associative learning in infants and adults to test the hypothesis that infants show superior crossmodal learning compared to adults when they encounter crossmodal associations passively. In contrast, adults learn crossmodal associations predominantly when task relevant. In the first experiment we tested a group of six-month-old infants (Experiment 1a) and a group of young adults (Experiment 1b). While recording EEG, we presented two frequently occurring audio-visual standard combinations (A1V1, A2V2, p = 0.35 each, ‘Frequent standard stimuli’), two rare recombinations of the standard stimuli (A1V2, A2V1, p = 0.10 each, ‘Rare recombined stimuli’) and one rare audio-visual combination of deviant auditory and deviant visual stimuli (A3V3, p = 0.10, ‘Rare deviant stimuli’). In a second experiment we tested an additional group of young adults in adapted versions of the same experiment: participants were not passively exposed to the stimuli, but had to respond to a target stimulus. In Experiment 2a participants had to detect a rare unimodal visual stimulus (V4) while the target stimulus in Experiment 2b was one of the rare recombined stimuli (A1V2 or A2V1). Thus, the crossmodal combinations were task relevant in Experiment 2b but not in Experiment 2a.

We predicted that infants would be able to discriminate between the frequent standard and rare deviant stimuli as well as between frequent standard and rare recombined stimuli, indicated by a deviant response in the event-related potentials (ERPs). Similar to the infant group we expected a deviant response to rare deviant stimuli in in all three experiments with adults. In contrast, a differentiation between standard and rare recombined stimuli was expected to emerge in adults only in Experiment 2b, that is when crossmodal combinations were task relevant.

## Methods

### Experiment 1

In Experiment 1 we investigated a group of infants (Experiment 1a) and a group of young adults (Experiment 1b) with the same experimental design. Due to the age difference between the groups adjustments in the procedure and data analyses were necessary. These are described below.

#### Participants: Experiment 1a

Sixty-two six-month-old infants (+/− 10 days) took part. Infants were recruited from the local registration offices. All participating infants were born full-term (38 – 41 weeks), had a typical prenatal and perinatal history and no known neurological or developmental problems. Parents gave their written consent and were informed about their right to abort the experiment at any time. They received a small present for their children (toy or picture book) for taking part. Thirty-three participants were excluded from the analyses because of too many artifacts in the EEG recordings, leaving a total of twenty-nine data sets for the final statistical analyses (17 female, 12 male). Note that an exclusion rate of approximately 50 % due to artifacts is not uncommon in infant research (DeBoer et al. 2007). Sample size of Experiment 1a and the following experiments was selected based on previous studies investigating typical sensory mismatch ERP effects. The study (including Experiment 1a and 1b) was performed in accordance with the ethical standards laid down in the Declaration of Helsinki in 1964. The procedure was approved by the ethics board of the German Psychological Society (DGPs).

#### Stimuli and Design: Experiment 1a

The experiment comprised three auditory and three visual stimuli, combined into crossmodal pairs of one visual and one auditory stimulus. All three auditory stimuli were presented with equal loudness but differed in sound frequency (400, 1000 or 1600 Hz); they were presented for 500 ms each via two loudspeakers. The visual stimuli consisted of three geometric shapes (circle, triangle, and square; size: 10°) combined with three different colors (green, red, and blue) and were presented in the middle of a computer screen for 500 ms.

Participants were exposed to two frequently occurring audio-visual standard combinations (A1V1, A2V2, each with p = 0.35, ‘Frequent standard stimuli’) and three infrequently occurring audio-visual deviant combinations. The latter consisted of (1) two rare recombinations of the auditory and visual stimuli comprising the standard stimuli (A1V2, A2V1, each with p = 0.10, ‘Rare recombined stimuli’) and (2) one rare audio-visual combination of a deviant auditory and a deviant visual stimulus (A3V3, p = 0.10, ‘Rare deviant stimuli’), not occurring in the combinations of the frequent standard stimuli and the recombined stimuli. The inter stimulus interval between the different crossmodal combinations amounted to 1500 ms. The types of crossmodal combinations and stimuli used as ‘Frequent standard stimuli’, ‘Rare recombined stimuli’, and ‘Rare deviant stimuli’ were counterbalanced over participants. The experiment was divided into five experimental blocks, each comprising 60 trials resulting in a total of 300 trials. For each block the proportion of the three conditions was 70: 20: 10 % (see Table 1). Thus, even if the experiment was prematurely aborted, each infant received the correct ratio of stimuli.

**Table 1.** Experimental design of Experiment 1a and Experiment 1b.

| Stimuli | Proportion | Condition (number of trials) |
| --- | --- | --- |
| Auditory 1 – Visual 1 (A1V1)<br>Auditory 2 – Visual 2 (A2V2) | 0.35 } 0.70<br>0.35 } | Frequent standard stimuli (210) |
| Auditory 1 – Visual 2 (A1V2)<br>Auditory 2 – Visual 1 (A2V2) | 0.10 } 0.20<br>0.10 } | Rare recombined stimuli (60) |
| Auditory 3 – Visual 3 (A3V3) | 0.10 } 0.10 | Rare deviant stimuli (30) |

#### Procedure: Experiment 1a

Experiment 1a took place in a sound-attenuated and electrically shielded room. During the experiment, the infants sat on their parents’ laps. The computer screen, displaying the visual stimuli, was positioned on a table at a distance of approximately 60 cm from the participants. Infants’ heads were aligned with the center of the screen. The two loud speakers were positioned behind the computer screen.

To make sure that the infants attentively observed the stimuli, a black and white video was continuously played in the background. This video consisted of 30 different sequences of centrally moving patterns, e.g. randomly moving stars or flying balloons focusing the viewing direction to the center of the computer screen. All sequences were ten seconds long and were presented without intermediate breaks. To control whether the infants were actually looking at the computer screen when the experimental visual stimuli were presented, a small camera, placed on top of the computer screen, recorded the infants’ heads. The camera was connected to the EEG recording computer to enable a continuous control of the child's attention as well as the EEG signal during the course of the experiment. If the infant did not look at the screen during the presentation of the stimuli, a marker was manually inserted by the experimenter in the EEG data file and the associated EEG segments were later taken out of the analysis. To avoid interfering signals, parents were instructed not to talk to their children during the time the EEG was recorded. Whenever the infant showed signs of discomfort or restlessness, the experiment was paused. Occasionally, a hand puppet was used during such breaks to keep the infants alert and to make sure that they attended to the computer screen when the experiment was continued. The EEG recording only continued if both the child and the parent were content. The testing time for all infants ranged between five and ten minutes (M = 7.2 minutes, SD=1.6). Together with the preparation time, the infants and their parents spent approximately forty-five minutes in the laboratory.

#### Electrophysiological recording and data analyses: Experiment 1a

EEG data were collected from 45 scalp sites using active Ag/AgCl electrodes (Brain Products, Easycap GmBH, Herrsching) mounted in an elastic cap (Electro Cap International, Inc.). The electrodes were placed according to the international 10-10 system (see Figure 1). EEG Data were recorded continuously using a band-pass filter of 0.01-250 with a sampling rate of 500 Hz (Brain Products, Munich, Germany). The electrode FPz served as online reference electrode and the ground electrode was applied at AF3. Data were re-referenced offline to the average of the recordings of electrodes TP9 and TP10, which are located close to the mastoids. Artifacts were rejected manually after visual inspection of each individual EEG trial. Trials with artifacts such as head movements, eye blinks, eye movements or electrical noises were removed from further analyses. The first 15 trials of each dataset were excluded since the participants were not yet familiarized with the relative proportions of each stimulus condition. Noisy channels were interpolated by calculating the average of the four adjacent electrodes (Picton et al., 2000). On average, three electrodes were interpolated for each participant. EEG data sets of infants (n=21) comprising less than 10 trials per condition were excluded from the final statistical analyses (see participants Experiment 1a).

For the statistical analyses, the lateral electrodes were grouped into four clusters for each hemisphere; each cluster comprised four electrodes (see Figure 1): the left hemisphere: (1) Frontal (F): F9, F7, F3, FC1; (2) Fronto-central (FC): FT9, FT7, FC5, C3; (3) Central-parietal (CP): T7, C5, TP7, CP5; (4) Parietal-occipital (PO): P3, P7, PO9, O1 and the right hemisphere: (1) Frontal (F): F10, F8, F4, FC2; (2) Fronto-central (FC): FT10, FT8, FC6, C4; (3) Central-parietal (CP): T8, C6, TP8, CP6; (4) Parietal-occipital (PO): P4, P8, PO10, O2. The midline electrodes AFz, Fz, FCz, Cz, Pz, and POz were separately analyzed. EEG data were segmented into epochs from 100 ms pre-stimulus to 1100 ms post-stimulus onset. Epochs were baseline corrected by means of the 100 ms pre-stimulus interval. The following two time windows were chosen based on a visual inspection of the group average ERPs: (1) 200 – 420 ms and (2) 420 – 1000 ms. To evaluate differences between conditions, a repeated measurement ANOVA comprising the within subject factors *Condition* (three levels: ‘Frequent standard stimuli’ vs. ‘Rare recombined stimuli’ vs. ‘Rare deviant stimuli’), *Hemisphere* (two levels: left vs. right) and *Cluster* (four levels: F vs. FC vs. CP vs. PO) was calculated separately for each of the two time windows.

**Figure 1.**
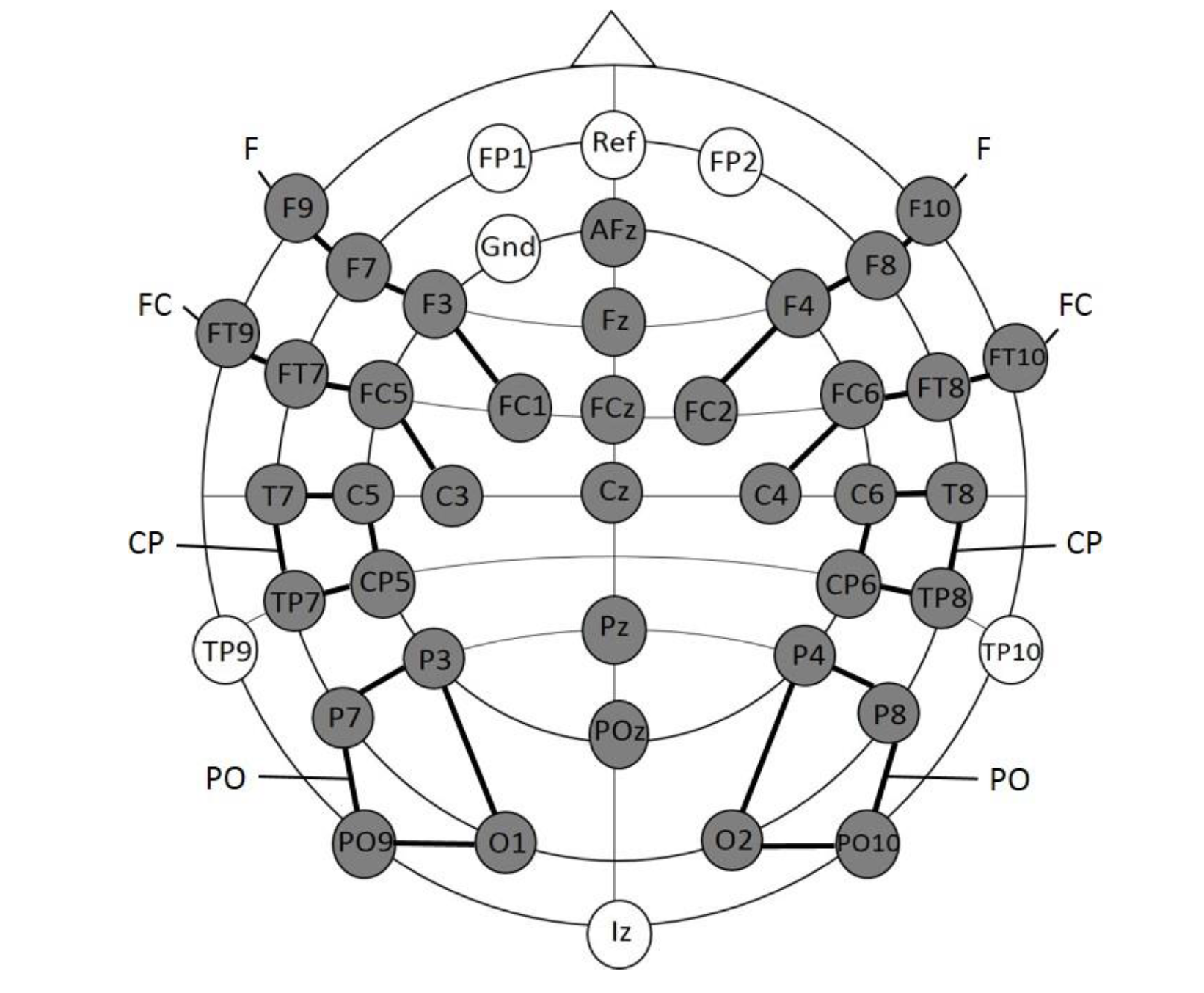
Electrode placement for experiment 1a; the grey electrodes were included in the statistical analyses. Clusters are indicated by black connecting lines and were named according to their location along the anterior-posterior axis.

Significant interactions including the factor *Condition* were followed up with sub-ANOVAs, calculated separately for each cluster. Significant main effects of *Condition* or interactions of *Condition* and *Hemisphere* were further analyzed with paired t-tests: 1) ‘Frequent standard stimuli’ vs. ‘Rare deviant stimuli’ and 2) ‘Frequent standard stimuli’ vs. ‘Rare recombined stimuli’. The midline electrodes were separately analyzed with an ANOVA comprising the factors *Condition* (three levels: Standard vs. New Combination vs. New Stimuli) and *Electrode* (six levels: AFz vs. Fz vs. Cz. vs. Pz vs. POz). Similar to the cluster analysis, significant interactions between the factor *Condition* and *Electrode* were further analyzed by calculating sub ANOVAs and paired t-tests separately for each electrode. The Huynh-Feldt correction was applied to all analyses comprising within subject factors with more than two levels. To correct for multiple comparisons, p-values of the t-tests were adjusted with the Bonferroni-Holm method. Only main effects and interactions, including the factor *Condition*, as well as significant post hoc tests are reported.

#### Participants: Experiment 1b

Twenty-seven young adults recruited from a student-subject database of the Institute for Psychology (University of Hamburg) were tested. They received either 8 €/ hour or course-credit. All participants had normal or corrected-to-normal vision, normal hearing and were free of neurological problems. All participants gave their informed consent. Four participants were excluded from the analysis due of too many artifacts in the EEG. A total of twenty-three participants were included in the final analyses (11 male, mean age 23.5 years, range 19-31)

#### Stimuli and Design: Experiment 1b

The stimuli and experimental design of Experiment 1b were identical to Experiment 1a (see Table 1).

#### Procedure: Experiment 1b

Experiment 1b took place in the adult EEG lab of the Biological Psychology and Neuropsychology section of the University of Hamburg. It was constructed by the same company as the Baby lab and had the same light sources, sound attenuating, and electrical shielding system. The experimental room was dimly lit and the participants were seated in a comfortable chair in front of a table. All devices used were the same as for Experiment 1a. The computer screen, displaying the visual stimuli and background video, was positioned at eye level on a table at a distance of approximately 60 cm from the participants (size of the visual stimuli: 7°). The two loud speakers were located behind the computer screen. Before the experiment started, participants received written instructions concerning the procedure of the experiment. In addition, they were asked to sit as still as possible, to limit their eye blinking during the recording of the experimental blocks and to continuously look at the fixation point. To control that the participants attended to the computer screen participants’ heads were recorded via a small camera, placed on top of the computer screen, during the experiment.

#### Electrophysiological recording and data analyses: Experiment 1b

EEG recording and data analyses were identical to Experiment 2a and 2b. Note, that the similar results for the ERPs to rare deviants in infants and adults, including the lateralization, exclude the possibility that differences in analyzing procedures contributed to the below reported other group differences.

## Experiment 2

In a second experiment we tested a group of additional young adults in two adapted versions of Experiment 1 (Experiment 2a and 2b). Experiment 2a and 2b differed in the employed target stimulus which had to be detected by the participants. The procedure and data analyses were the same for both experiments.

### Participants

Seventeen healthy university students took part in the experiment. The participants were recruited from a student-subject database of the Institute of Psychology at the University of Hamburg. They received either 8 €/ hour or course-credit. All participants had normal or corrected-to-normal vision, normal hearing and no neurological problems. Five participants were excluded from the analysis due to too many artifacts in the EEG or insufficient task performance (less than 70 % correct target detection), leaving a total of twelve participants for the final analyses (four male, age 20 – 31 years, mean = 23.8 years). All participants gave their informed consent. The study was performed in accordance with the ethical standards laid down in the Declaration of Helsinki in 1964. The procedure was approved by the ethics board of the German Psychological Society (DGPs).

### Stimuli and design

The design of Experiment 2 was similar to Experiment 1, but the stimuli and the experimental setting was adjusted. A visual LED was located inside a small wooden front (22 × 24 cm) which was covered with a black cloth. The wooden front was placed on top of a black box, to make sure that the position of the LED was at eye-level at a distance of approximately 85 cm from the participants. The LED was activated for 100 ms in four possible colors: red, blue, green or yellow. Auditory stimuli (400, 800, or 1600 Hz) were presented for 100 ms via two speakers which were positioned adjacent to the wooden front. Crossmodal stimuli were made by combining one of the sounds with one of the LED colors. Crossmodal combinations were counterbalanced over conditions and participants. In contrast to Experiment 1b, adults were engaged in a task and had to detect a target stimulus rather than being passively exposed to a sequence of crossmodal stimuli. The target stimulus was either unrelated to the crossmodal combinations (Experiment 2a) or addressed specific crossmodal combinations (Experiment 2b), resulting in two different experiments.

In Experiment 2a the frequent standard stimuli (A1V1, A2V2) were presented with a probability of p = 0.30 each while the rare recombined (A1V2, A2V1) and rare deviant stimuli (A3V3) had a probability of p = 0.10 each. An additional unimodal visual stimulus (p = 0.10, V4) served as target stimulus (see Table 2A).

**Table 2.**
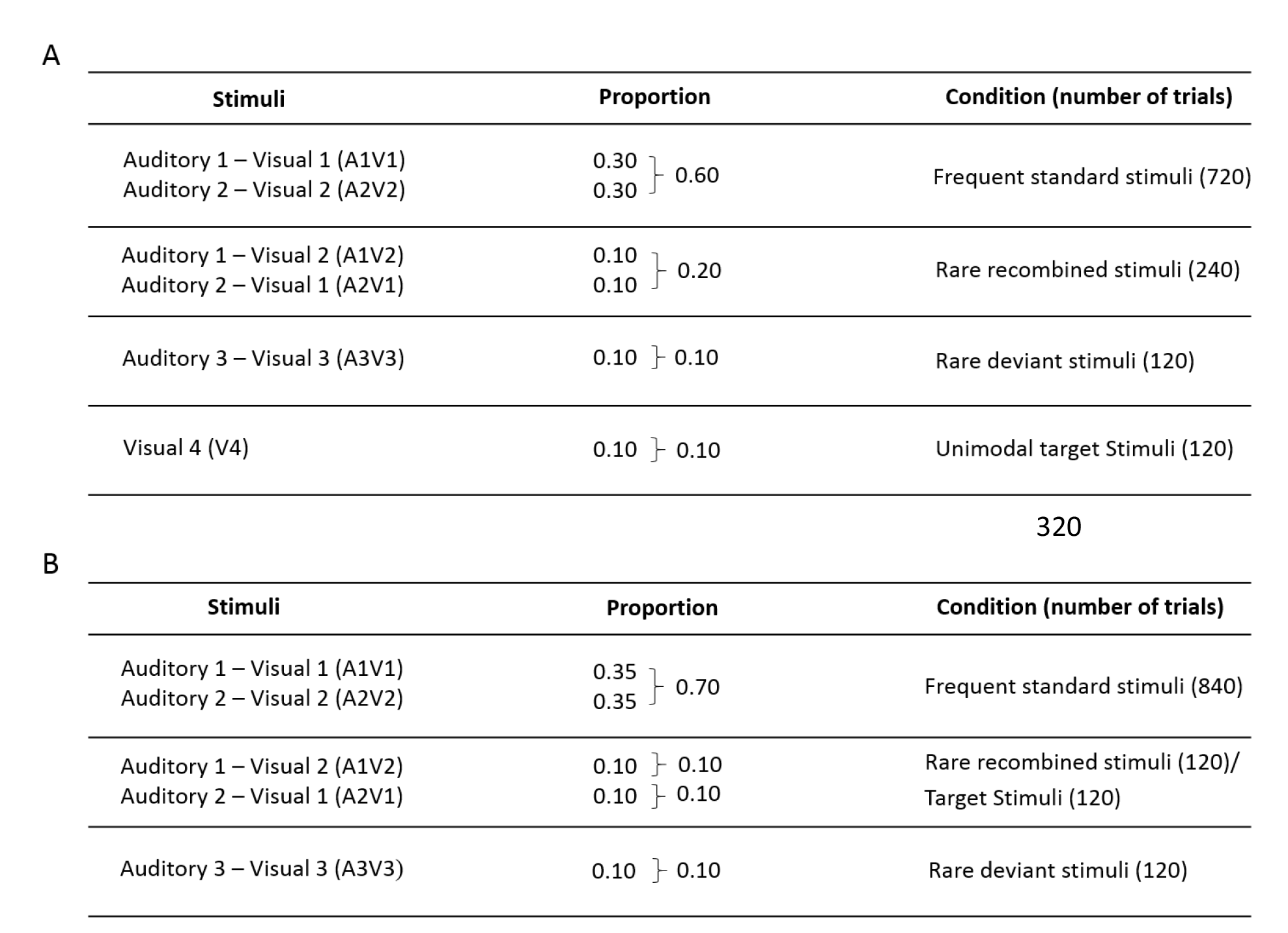
Experimental design of A) Experiment 2a and B) Experiment 2b.

| Stimuli | Proportion | Condition (number of trials) |
| --- | --- | --- |
| Auditory 1 – Visual 1 (A1V1)<br>Auditory 2 – Visual 2 (A2V2) | 0.30 } 0.60<br>0.30 } | Frequent standard stimuli (720) |
| Auditory 1 – Visual 2 (A1V2)<br>Auditory 2 – Visual 1 (A2V1) | 0.10 } 0.20<br>0.10 } | Rare recombined stimuli (240) |
| Auditory 3 – Visual 3 (A3V3) | 0.10 } 0.10 | Rare deviant stimuli (120) |
| Visual 4 (V4) | 0.10 } 0.10 | Unimodal target Stimuli (120) |

B
| Stimuli | Proportion | Condition (number of trials) |
| --- | --- | --- |
| Auditory 1 – Visual 1 (A1V1)<br>Auditory 2 – Visual 2 (A2V2) | 0.35 } 0.70<br>0.35 } | Frequent standard stimuli (840) |
| Auditory 1 – Visual 2 (A1V2)<br>Auditory 2 – Visual 1 (A2V1) | 0.10 } 0.10<br>0.10 } 0.10 | Rare recombined stimuli (120)/<br>Target Stimuli (120) |
| Auditory 3 – Visual 3 (A3V3) | 0.10 } 0.10 | Rare deviant stimuli (120) |

In Experiment 2b there was no unimodal V4, but the target stimulus was defined as one of the rare recombined stimuli (either A1V2 or A2V1) rendering crossmodal combinations task relevant. A1V1 and A2V2 were presented with a probability of p = 0.35 each while the probability for A1V2, A2V1, and A3V3 was p = 0.10 each (see Table 2B). All participants took part in both experiments. The order of the two experiments as well as the specific audio-visual combinations used for the different conditions were counterbalanced over participants. Stimuli were presented in six blocks with 200 trials per block.

### Procedure

The experiment took place in a dimly lit, sound-attenuating, and electrical shielded room. The participants were seated in a comfortable chair at a table approximately 85 cm from the box that contained the visual LED. The target stimulus was presented three times prior to the start of the experiment, to allow participants to get acquainted with the target. Responses to the target stimuli were made by means of a custom made button box, placed near the dominant hand. Participants were instructed to sit as still as possible and to keep their eyes focused on the LED. Experiment 2a and 2b lasted for twenty to thirty minutes each (including breaks). The total testing time, which included briefing of the participant, practice trails and EEG application, was approximately 1 hour and 45 minutes for both experiments.

### Behavioral analysis

All button presses within 100 and 1000 ms following stimulus presentation were considered valid responses. Hit, miss and false alarm rates were calculated and average reaction times to targets were derived for both Experiment 2a and 2b.

### Electrophysiological recording and data analysis

EEG data were collected from 74 scalp sites using active Ag/AgCl electrodes (Brain Products, Easycap GmBH, Herrsching) mounted on an elastic cap (Electro Cap International, Inc.). Data were recorded continuously using a band-pass filter of 0.01-250 with a sampling rate of 500 Hz (Brain Products, Munich, Germany). The electrodes were placed according to the international 10-10 system (see Figure 2). One additional electrode was positioned below the left eye to record vertical eye movements. A left earlobe electrode served as online reference electrode. EEG data were filtered offline with a low-pass filter with a 40 Hz cut-off and were re-referenced offline to an average reference. Electrodes positioned close to the outer canthi of each eye (F9 and F10) served for recording horizontal eye movements. An independent component analysis (ICA) was run for each EEG data set, which defined 30 time-independent components representing the data (Makeig, Debener, Onton & Delorme, 2004). Components representing artifacts such as eye blinks, eye movements, electrical noise or heart beat were manually detected and rejected from further analyses. The first 75 trials (Experiment 2a and 2b) or the first 15 trials (Experiment 1b) of each dataset were excluded since the participants were not yet familiarized with the relative proportions of each stimulus condition. The lateral electrodes were grouped into six clusters for each hemisphere; each cluster comprised five electrodes (see Figure 2): (1) Frontal (F): F1, F3, F5, F7, F9 (2) Fronto-central (FC): FC1, FC3, FC5, FT7, FT9 (3) Central (C): C1, C3, C5, T7 (4) Centro-parietal (CP): CP1, CP3, CP5, TP7, TP9 (5) Parietal (P): P1, P3, P5, P7, P9 (6) Parieto-occipital (PO): PO3, PO7, PO9, O1, O9) and for the right hemisphere: (1) Frontal (F): F2, F4, F6, F8, F10 (2) Fronto-central (FC): FC2, FC4, FC6, FT8, FT10 (3) Central(C): C2, C4, C6, T8 (4) Centro-parietal (CP): CP2, CP4, CP6, TP8, TP10 (5) Parietal (P): P2, P4, P6, P8, P10 (6) Parieto-occipital (PO): PO4, PO8, PO10, O2, O10). The midline electrodes Fz, FCz, Cz, CPz, Pz, POz, and Oz were separately analyzed. EEG data were segmented into epochs starting 100 ms before the stimulus onset and lasting for 1000 ms post stimulus onset. Epochs were baseline corrected with a pre-stimulus interval of 100 ms. The following time epochs were chosen based on visual inspection of the group mean average: Experiment 2a: (1) 80 – 190 ms and (2) 250 – 850 ms; Experiment 2b (1) 80 – 160 ms, (2) 170 – 230 ms and (3) 250 – 850 ms. The statistical analyses were the same as described for Experiment 1a.

**Figure 2.**
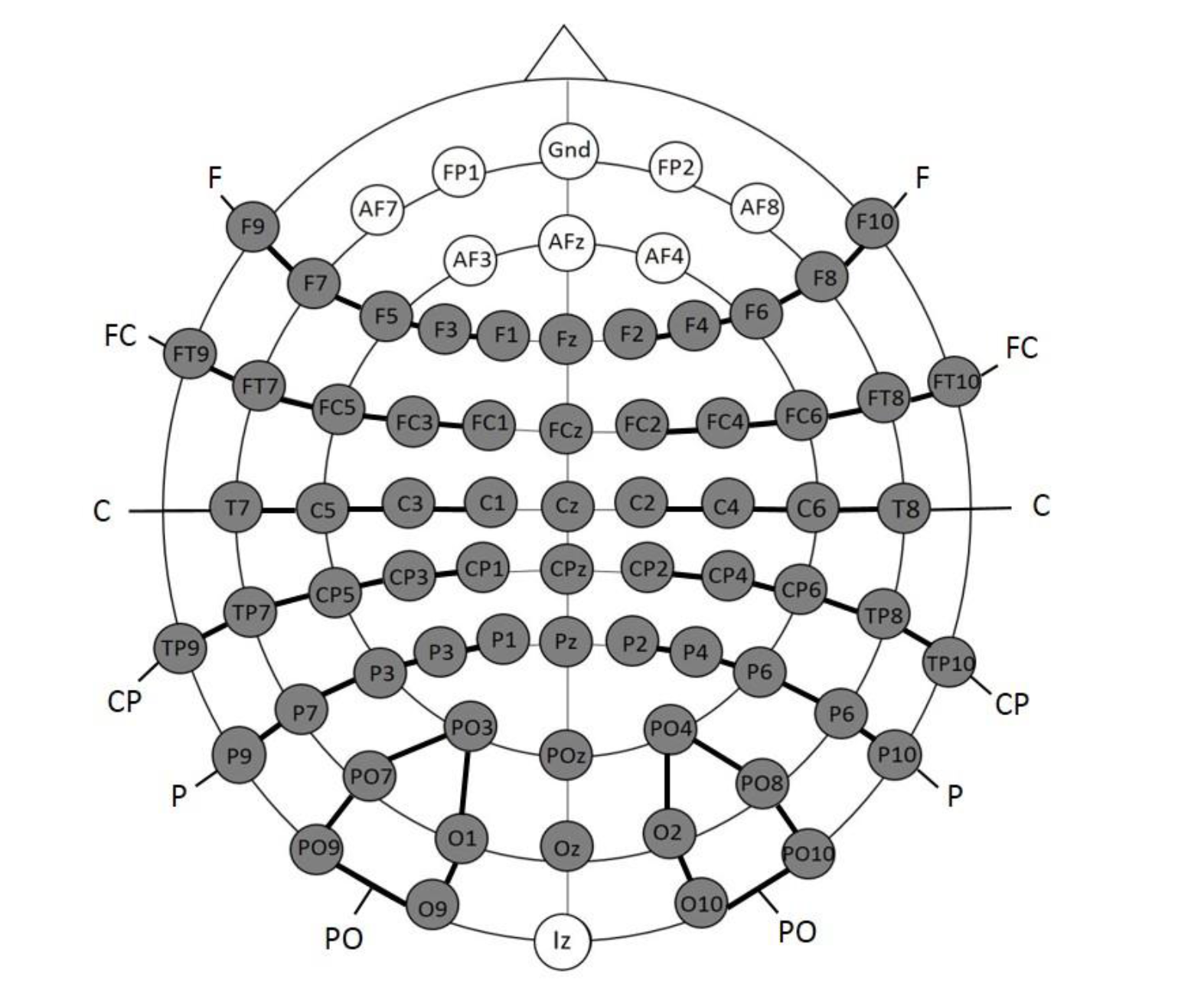
Electrode placement for Experiment 2a and 2b; the grey electrodes were included in the statistical analyses. Clusters are indicated by black connecting lines and were named according to their location along the anterior-posterior axis.

## Results

### Experiment 1a (Infants)

Rare deviant stimuli (A3V3) elicited a more negative going ERP than audio-visual standard stimuli (A1V2, A2V2) (see Figure 3). This effect (200-420 ms, 420-1000 ms) was predominantly observed over the right hemisphere. Crucially, rare recombined stimuli (A1V2, A2V2) elicited a more positive going ERP compared to frequent standards (see Figure 3), predominantly over the left hemisphere (420 – 1000 ms).

#### First time window (200 – 420 ms) cluster analysis

The overall ANOVA with factors *Condition, Hemisphere*, and *Cluster* revealed a significant interaction between the factors *Condition* and *Hemisphere* (F(2,56) = 4.55; *P* = 0.015) as well as a significant interaction of *Condition × Cluster* (F(6,168) = 4.94; *P* < 0.001). Follow-up ANOVAs revealed a significant interaction of *Condition* × *Hemisphere* for cluster F (F(2,56) = 3.78; *P* = 0.028), FC (F(2,56) = 3.67; *P* = 0.029), and cluster CP (F(2,56) = 3.18; *P* = 0.048). Post hoc t-tests showed that this interaction was driven by a more positive amplitude in response to rare deviant stimuli compared to standard stimuli (see Figure 3) at cluster F (t(28) = 3.18; P = 0.014), cluster FC (t(28) = 2.93; *P* = 0.026), and cluster CP (t(28) = 3.02; *P* = 0.02) of the right hemisphere.

#### First time window (200 – 420 ms) midline analysis

The overall ANOVA with factors *Condition* and *Electrode* showed a significant interaction between *Condition* x *Electrode* (F(10,280) = 2.76; *P* = 0.002). Follow-up ANOVAs revealed a significant main effect of the factor *Condition* for electrode Fz (F(2,56) = 5.3; *P* = 0.007) and FCz (F(2,56) = 3.79; *P* = 0.02). Post hoc t-tests showed significant differences between the rare deviant and standard condition at electrode FC (t(28) = 2.5; *P* = 0.036) and FCz (t(28) = 2.45; *P* = 0.04); rare deviant stimuli elicited a more positive going ERP than standard stimuli (see Figure 3).

**Figure 3.**
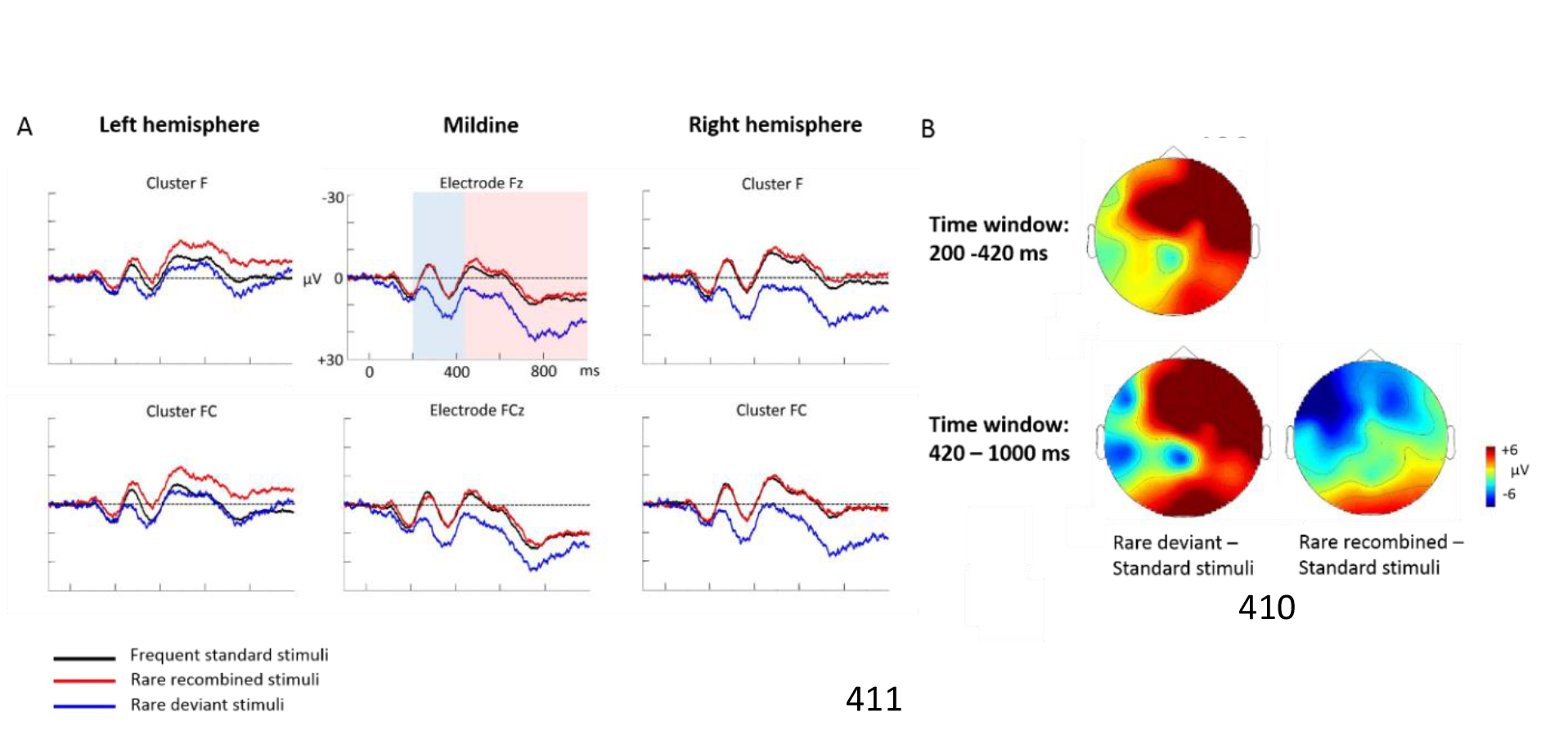
Grand average ERPs of Experiment 1a. A) ERPs to the three conditions (frequent standard stimuli, rare recombined stimuli, rare deviant stimuli) are superimposed for the electrode clusters F and FC, and the electrodes Fz and FCZ. The analyzed time epochs are marked in blue (200-420 ms) and red (420-1000 ms). B) The topographical distribution of the difference between ‘Standard stimuli’- ‘Rare deviant stimuli" and ‘Standard’- ‘Rare recombined stimuli’ for the first and second time window.

#### Second time window (420 – 1000 ms) cluster analysis

The overall ANOVA revealed a significant interaction of *Condition* × *Hemisphere* (F(2,56) = 4.68; *P* = 0.013) as well as a significant interaction of *Condition × Cluster* (F(6,168) = 4.51; *P* < 0.01). Follow-up ANOVAs showed a significant interaction of *Condition × Hemisphere* at Cluster F (F(2,56) = 4.5; *P* = 0.014) and cluster FC (F(2,56) = 4.6; *P* = 0.013). Post-hoc t-tests indicated that ERPs to rare deviant stimuli were significantly more positive than ERPs to standard stimuli (see Figure 3) at cluster F (t(28) = 2.72; *P* = 0.044) of the right hemisphere. In addition, post hoc t-tests revealed significant differences between standard and rare recombined stimuli at cluster FC of the left hemisphere (t(28) = −2.81; *P* = 0.032), indicating a more negative amplitude in response to rare recombined stimuli compared to the standard stimuli (see Figure 3).

Second time window (420 – 1000 ms): midline analysis. The ANOVA revealed a significant interaction between the factors *Condition* and *Electrode* (F(10,280) = 2.76; *P* = 0.002). Follow-up ANOVAs indicated a main effect of *Condition* for electrode AFz (F(2,56) = 3.4; *P* = 0.04) and Fz (F(2,56)= 3.59; *P* = 0.03). However, none of the subsequent t-tests reached significance (all p > 0.08).

### Experiment 1b (Adults)

ERPs to rare deviant stimuli were more negative going than ERPs to standard stimuli during both time windows (180-220 ms, 250 −1000 ms; see Figure 4).

**Figure 4:**
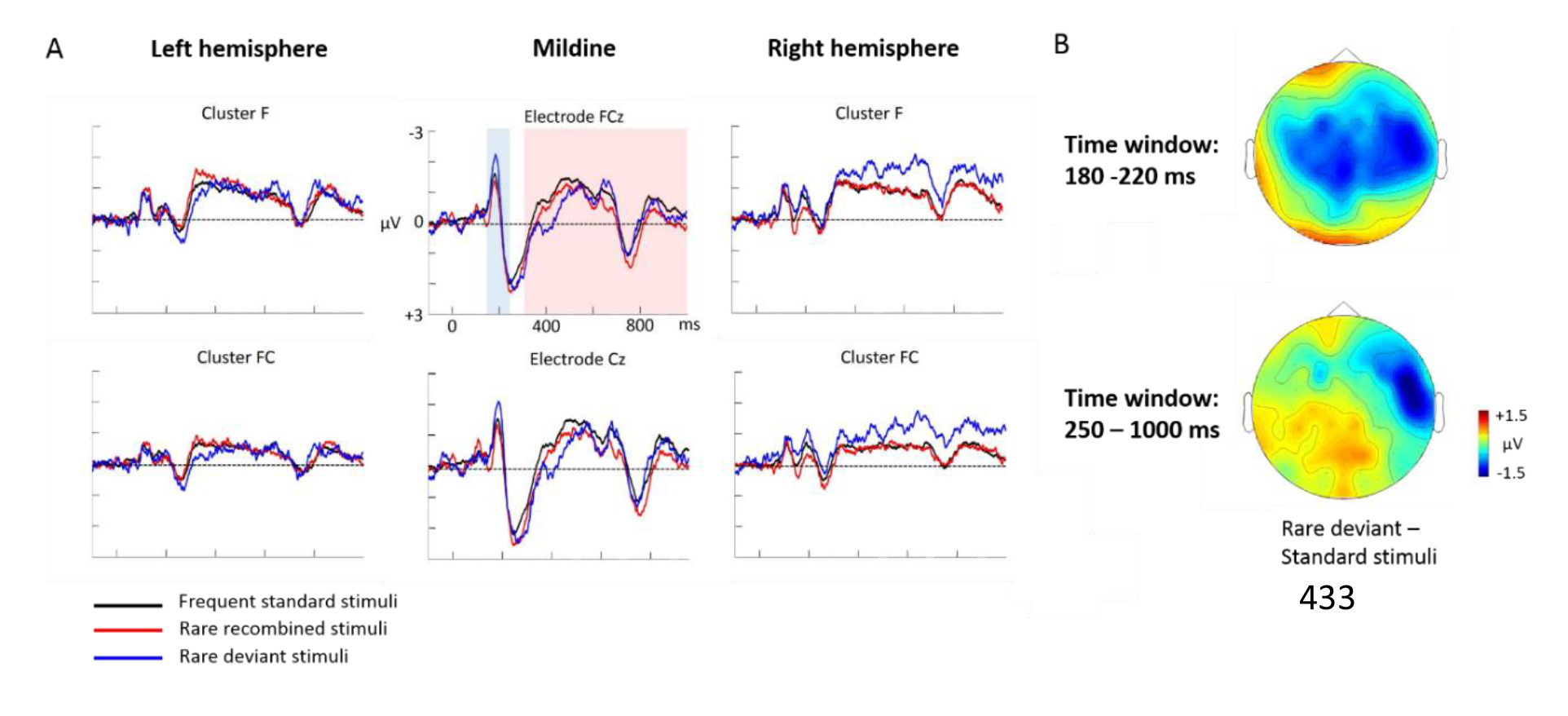
Grand average ERPs of Experiment 1b. A) ERPs to the three conditions (frequent standard stimuli, rare recombined stimuli, rare deviant stimuli) are superimposed for the electrode clusters F and FC, and the electrodes FCz and Cz. The analyzed time epochs are marked in blue (180-220 ms) and red (420-1000 ms). B) The topographical distribution of the difference between ‘Standard stimuli’- ‘Rare deviant stimuli’ for the first and second time window.

#### First time window (180 -220 ms) cluster analysis

The overall ANOVA did not reveal any significant effect involving the factor *Condition*.

#### First time window (180 -220 ms) midline analysis

The overall ANOVA revealed a significant interaction between the factors *Condition* and *Electrode* (F(12,276) = 2.16; *P* = 0.03). Follow-up ANOVAs revealed a significant main effect of *Condition* for electrode CPz (F(2,46) = 4.02; *P* = 0.024). Post hoc t-tests showed significant differences between the rare deviant and standard stimuli at electrode Cz (t(22) = 2.32; *P* = 0.047); rare deviants elicited a more negative going ERP than standard stimuli (see Figure 4).

#### Second time window (250 – 1000 ms) cluster analysis

The overall ANOVA revealed a significant interaction between the factors *Condition, Hemisphere*, and *Cluster* (F(10,230) = 2.49; *P* = 0.007). Follow-up ANOVAs obtained a significant main effect of *Condition* for cluster FC (F(2,46) = 4.56; *P* = 0.015). Post hoc t-tests showed that this interaction was driven by a more positive amplitude in response to rare deviant stimuli compared to standard stimuli (see Figure 4) at cluster FC (t(22) = 2.22; *P* = 0.036).

#### Second time window (250 – 1000 ms) midline electrodes

The overall ANOVA did not reveal any significant effect involving factor *Condition*.

### Experiment 2: Behavioral data

As seen in Table 3, participants identified target stimuli with a high accuracy in both experiments.

**Table 3.**
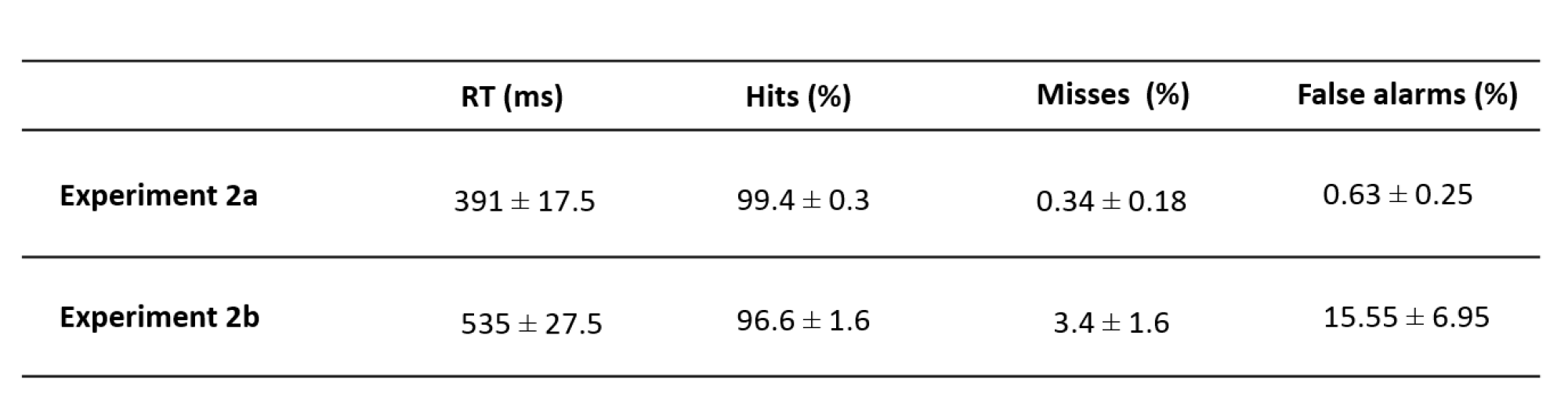
Mean (± SEM) of reaction time (in ms), hit rates (in %), misses (in %), and false alarms (in %) to the target stimuli of Experiment 2a and Experiment 2b.

|  | RT (ms) | Hits (%) | Misses (%) | False alarms (%) |
| --- | --- | --- | --- | --- |
| **Experiment 2a** | 391 ± 17.5 | 99.4 ± 0.3 | 0.34 ± 0.18 | 0.63 ± 0.25 |
| **Experiment 2b** | 535 ± 27.5 | 96.6 ± 1.6 | 3.4 ± 1.6 | 15.55 ± 6.95 |

### Experiment 2a: ERP data

Rare deviant stimuli elicited more negative going ERPs compared to standard stimuli (80-190 ms and 250-850 ms) while ERPs to standard and rare recombined stimuli did not significantly differ (see Figure 5).

**Figure 5.**
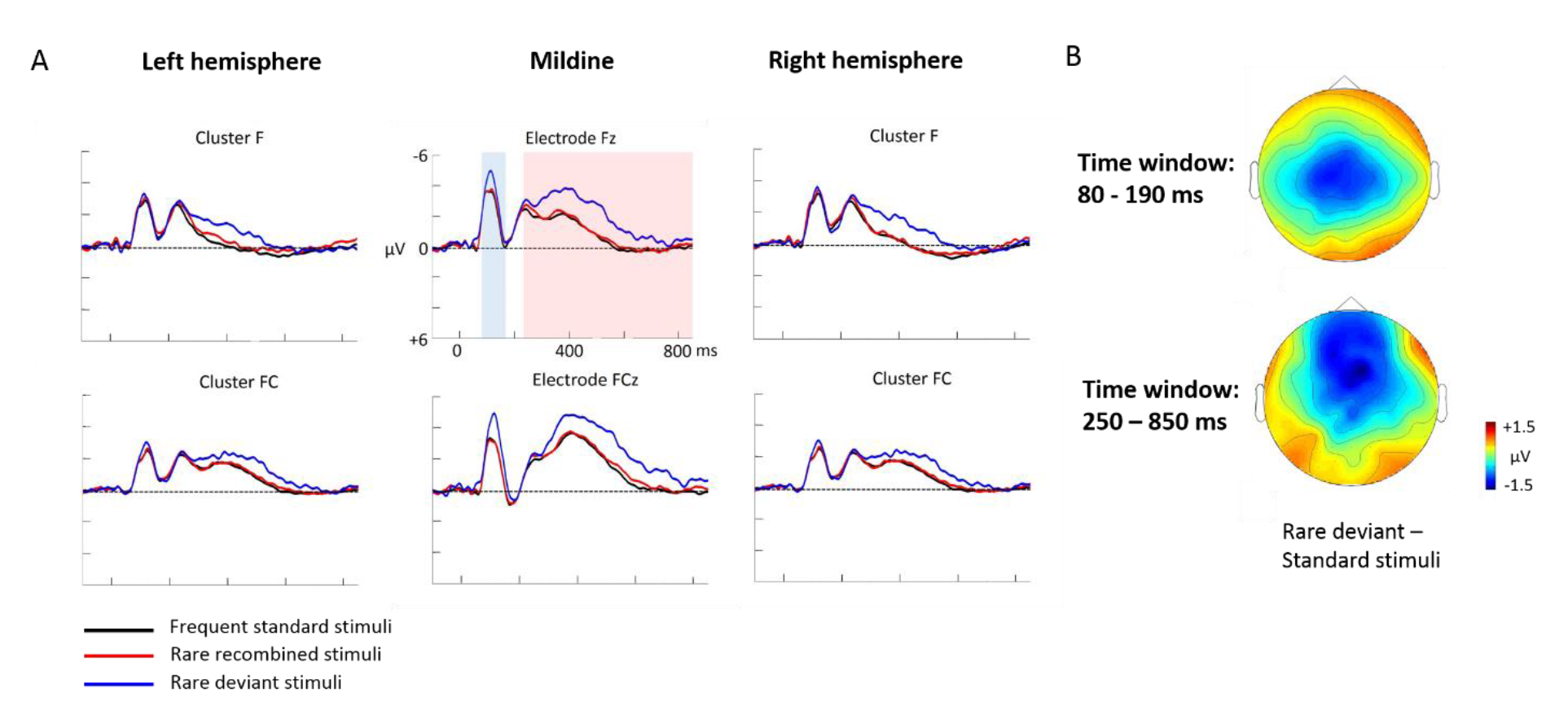
Grand average ERPs of Experiment 2a. A) ERPs to the three conditions (frequent standard stimuli, rare recombined stimuli, rare deviant stimuli) are superimposed for the electrode clusters F and FC, and the electrodes Fz and FCZ. The analyzed time epochs are marked in blue (80-190 ms) and red (250-850 ms). B) The topographical distribution of the difference between ‘Standard stimuli’- ‘Rare deviant stimuli" for the first and second time window.

#### First time window (80 – 190 ms) cluster analysis

The overall ANOVA revealed a significant interaction between the factors *Condition* and *Cluster* (F(10,110) = 2.74; *P* < 0.001). Follow-up ANOVAS showed a significant main effect of *Condition* for cluster C (F(2,22) = 18.85; *P* < 0.001) and cluster CP (F(2,22) = 3.84; *P* = 0.034). Post-hoc t-tests indicated that ERPs to rare deviant stimuli were significantly more negative than ERPs to standard stimuli (see Figure 5) at cluster C (t(11) = 4.93; *P* < 0.001).

#### First time window (80 – 190 ms) midline analysis

The overall ANOVA revealed a significant interaction of *Condition* × *Electrode* (F(12,132) = 3.76; *P* < 0.001). Follow-up ANOVAs obtained a significant main effect of *Condition* for electrode FCz (F(2,22) = 11.88; *P* < 0.001), Cz (F(2,22) = 15.34; *P* < 0.001), CPz (F(2,22) = 20.44; *P* < 0.001), Pz (F(2,22) = 10.48; *P* < 0.001). Subsequent t-tests showed that this main effect was driven by a significant more negative amplitude in response to the rare deviant stimuli compared to the standard stimuli (see Figure 5) at electrode FCz (t(11) = −3.99; *P* = 0.003), Cz (t(11) = −4.48; *P* = 0.001), CPz (t(11) = −4.86; *P* < 0.001), and Pz (t(11) = −2.81; *P* = 0.029).

#### Second time window (250 – 850 ms) cluster analysis

The overall ANOVA revealed an interaction between *Condition* × *Cluster* (F(10,110) = 3.23; *P* < 0.001). Follow-up ANOVAs showed a significant main effect of factor *Condition* for cluster P (F(2,22) = 4.9; *P* = 0.015) and cluster PO (F(2,22) = 4.74; *P* = 0.017). Post-hoc t-tests indicated that ERPs in response to rare deviant stimuli were significantly more negative compared to ERPs to standard stimuli (see Figure 5) at cluster P (t(11) = 3.46; *P* = 0.008) and cluster PO (t(11) = 3.47; *P* = 0.008)

#### Second time window (250 – 850 ms) midline analysis

The overall ANOVA revealed a significant interaction of *Condition* × *Electrode* (F(12,132) = 3.82; *P* < 0.001). Sub ANOVAs showed a significant main effect for the factor *Condition* at electrode Fz (2,22) = 10.59; *P* < 0.001), FCz (F(2,22)= 8.86; *P* = 0.001), Cz (F(2,22) = 4.13; *P* = 0.027). Subsequent t-tests detected significant differences between the standard and rare deviant stimuli at electrode Fz (t(11) = 5.71; *P* < 0.001), FCz (t(11) = 4.49; *P* = 0.001), and Cz (t(11) = 2.53; *P* = 0.049); ERPs to rare deviants were more negative going than ERPs to standard stimuli (see Figure 5).

### Experiment 2b: ERP data

ERPs to rare deviant stimuli were more negative going than ERPs to standard stimuli (80-160 ms, 170-230 ms, 250-850 ms). Crucially, ERPs to rare recombined stimuli were more positive going than to standards (250-850 ms; see Figure 6).

**Figure 6.**
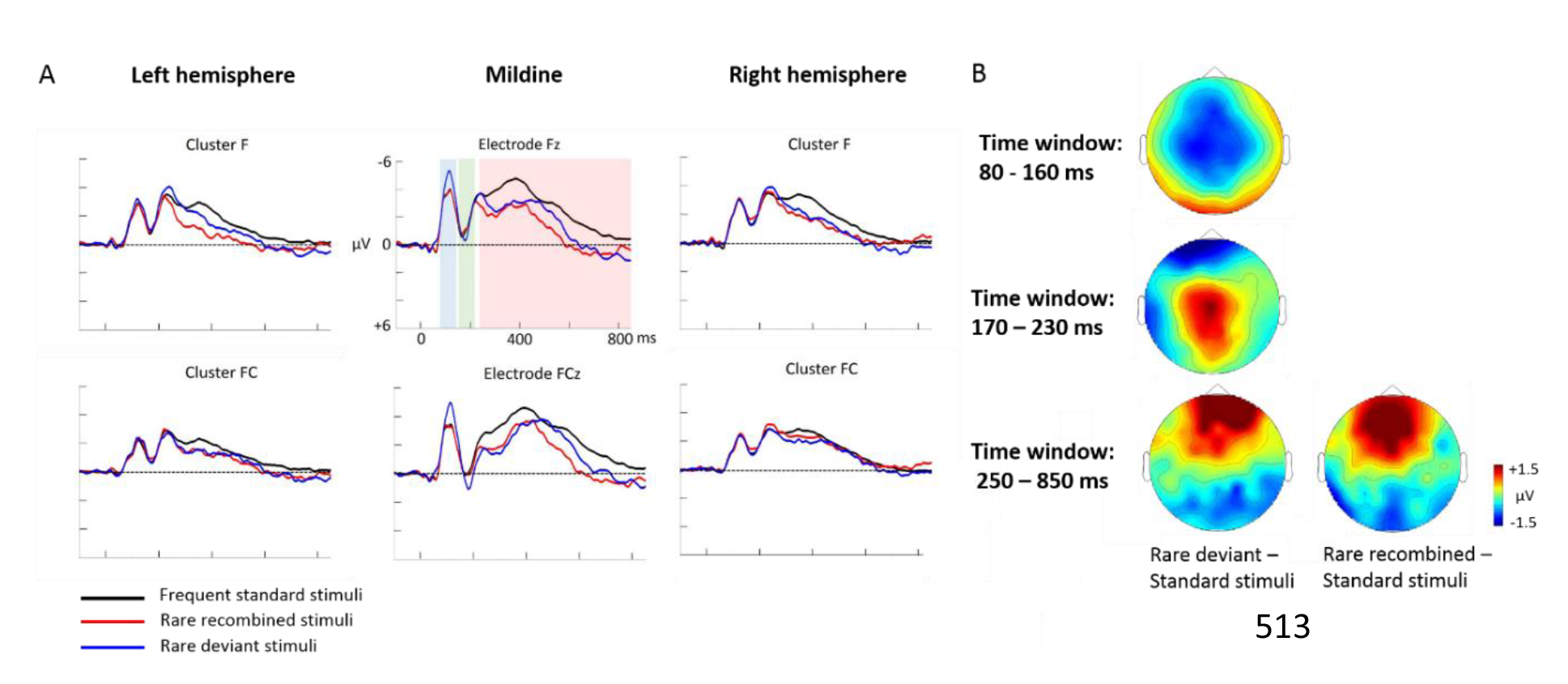
Grand average ERPs of Experiment 2b. A) ERPs to the three conditions (frequent standard stimuli, rare recombined stimuli, rare deviant stimuli) are superimposed for the electrode clusters F and FC, and the electrodes Fz and FCZ. The analyzed time epochs are marked in blue (80-160 ms), green (170-230 ms) and red (250-850 ms). B) The topographical distribution of the difference between ‘Standard stimuli’- ‘Rare deviant stimuli’ and ‘Standard stimuli’ - ‘Rare recombined stimuli’ for the first and second time window.

#### First time window (80 – 160 ms) cluster analysis

The overall ANOVA revealed a significant interaction of *Condition* × *Cluster* (F(10,110) = 3.82; *P* = 0.044). Further sub-ANOVAs showed a main effect of *Condition* for cluster C (F(2,22) = 5.83; *P* = 0.003) and cluster PO (F(2,22) = 4.16; *P* = 0.027), indicating a significant more negative amplitude in response to rare deviant than to standard stimuli (see Figure 6) at cluster C (t(11) = 4.44; *P* = 0.001) and cluster PO (t(11) = 3.19; *P* = 0.014).

#### First time window (80 – 160 ms) midline analysis

The overall ANOVA revealed a significant interaction of *Condition* × *Electrode* (F(12,132) = 2.72; *P* = 0.002). Follow-up ANOVAs revealed a significant main effect of *Condition* for electrode FCz (F(2,22) = 4.28; *P* = 0.024), Cz (F(2,22) = 6.01; *P* = 0.007) and CPz (F(2,22) = 3.67; *P* = 0.039). Subsequent t-tests indicated that ERPs to rare deviant were more negative than to standard stimuli (see Figure 6) at electrode FCz (t(11) = −2.85; *P* = 0.026), Cz (t(11) = −3.59; *P* = 0.006), and CPz (t(11) = −2.59; *P* = 0.044).

#### Second time window (170 – 230 ms): cluster analysis

The overall ANOVA did not reveal any significant effect involving factor *Condition*.

#### Second time window (170 – 230 ms): midline analysis

The overall ANOVA showed a significant interaction of *Condition* × *Electrode* (F(12,132) = 4.16; *P* = 0.01). Follow-up ANOVAs revealed a main effect of *Condition* for electrode FCz (F(2,22) = 3.44; *P* = 0.047), Cz (F(2,22) = 7.17; *P* = 0.003), CPz (F(2,22) = 11.47; *P* < 0.001), and Pz (F(2,22) = 20.37; *P* < 0.001). Subsequent t-tests indicated more positive going ERPs to rare deviant than to standard stimuli (see Figure 6) at electrode FCz (t(11) = 3.05; *P* = 0.018), Cz (t(11) = 3.74; *P* = 0.005), CPz (t(11) = 3.87; *P* = 0.003), and Pz (t(11) = 3.7; *P* = 0.005).

#### Third time window (250 −850 ms): cluster analysis

The overall ANOVA revealed a significant interaction of *Condition* × *Cluster* (F(10,110) = 4.12; *P* < 0.001). Follow-up ANOVAs showed a significant main effect of *Condition* for cluster F (F (2,22) = 5.09; *P* = 0.013), FC (F(2,22) = 4.4; *P* = 0.022), CP (F(2,22) = 6.42; *P* = 0.005), and PO (F(2,22) = 6.35; *P* = 0.005). Subsequent t-tests indicated significant more positive going ERPS to rare deviant than to standard stimuli (see Figure 6) at cluster F (t(11) = 2.77; *P* = 0.03), FC (t(11) = 3.88; *P* = 0.004), CP (t(11) = 2.62; *P* = 0.041), and PO (t(11) = 3.6; *P* = 0.01). In addition, t-tests showed that ERPs to rare recombined standards were more positive going than to standard stimuli (see Figure 6) at cluster F (t(11) = −3.11; *P* = 0.016), CP (t(11) = −3.43; *P* = 0.009), and PO (t(11) = −3.41; *P* = 0.016).

#### Third time window (250 −850 ms): midline analysis

The overall ANOVA revealed a significant interaction between *Condition* × *Electrode* (F(12,132) = 7.62; *P* < 0.001). Follow-up ANOVAs showed a main effect of *Condition* for electrode Fz (F(2,22) = 7.42; *P* = 0.003), FCz (F(2,22) = 9.24; *P* < 0.001), Cz (F(2,22) = 9.24; *P* < 0.001), Pz (F(2,22) = 6.49; *P* = 0.005), POz (F(2,22) = 7.92; *P* = 0.002), and Oz (F(2,22) = 5.62; *P* = 0.009). Subsequent t-tests indicated that ERPs to rare deviants were more negative going than to standard stimuli (see Figure 6) at electrode Fz (t(11) = 2.86; *P* = 0.013), FCz (t(11) = 3.71; *P* = 0.002), Pz (t(11) = 3.23; *P* = 0.006), POz (t(11) = 2.93; *P* = 0.01), and Oz (t(11) = −2.54; *P* = 0.024). Additionally, t-tests confirmed more positive going ERPs to rare recombined than to standard stimuli (see Figure 6) at electrode Fz (t(11) = −3.54; *P* = 0.01), FCz (t(11) = −4.29; *P* = 0.002), Pz (t(11) = −3.49; *P* = 0.003), POz (t(11) = −3.58; *P* = 0.006), and Oz (t(11) = −3.29; *P* = 0.01).

## Discussion

The goal of the present study was to test for a higher sensitivity of infants as compared to adults to crossmodal statistics and to compare the mechanisms of crossmodal association learning in infants and adults. We conducted ERP studies in which infants and adults were exposed to audio-visual stimulus combinations with different probabilities. ERPs to standard crossmodal combinations with a high frequency and to rare recombinations of these standards were compared. While infants passively learned the crossmodal combinations, adults discriminated recombined from standard combinations only when they were task relevant. In contrast, all groups succeeded in differentiating high frequent standard stimuli from rare audio-visual stimuli, which comprised infrequent auditory and visual elements.

Studies using artificial languages or visual artificial scenes have repeatedly demonstrated that infants develop a sensitivity to the likelihood of events as well as to conditional probabilities (Krogh et al., 2013; Aslin, 2014). Two recent studies found that six-month and twelve-month-old infants were able to learn to predict a visual stimulus based on a co-occurring or preceding auditory stimulus (Emberson et al., 2011; Kouider et al., 2015). While Kouider et al. (2015) demonstrated that infants at the age of twelve months were able to learn an association between an arbitrary sound and a visual object category (faces vs. flowers), they did not include an adult control group to demonstrate differences in learning between adults and infants, nor were they able to distinguish processes related to the detection of crossmodal combinations and the familiarity with certain sensory elements.

Thus, the present study extended previous research by showing that the probabilities of crossmodal combinations were extracted by infants as young as six months after a short exposure period while adults failed to learn crossmodal statistics under this condition. It is important to notice that we controlled for the likelihood of the auditory and visual elements of the employed crossmodal stimuli by recombining the auditory and visual elements of the frequent standard combinations. We provide ERP evidence demonstrating that the processing of crossmodal combinations and the processing of the likelihood of sensory elements can be dissociated: in infants, rare recombined stimuli elicited a left negative potential starting at about 420 ms post-stimulus while rare deviant stimuli elicited right lateralized positivity starting at 200 ms post-stimulus (Experiment 1a). Adults tested under identical conditions were only able to distinguish between rare deviant and standard stimuli (Experiment 1b, ERP effect starting 180 ms post-stimulus) but not between standard and rare recombined stimuli. These results demonstrate that infants were able to learn arbitrary crossmodal associations as early as six months of age and thus much earlier than suggested by the study of Kouider et al. (2015). Moreover, we provide first evidence that the learning of crossmodal statistics at this age is particularly sensitive and superior to adults. It could be argued that the signal to noise ratio of the ERPs in adults was not sufficient to demonstrate crossmodal learning in this group. However, two findings render this account for the present results unlikely: first, adults showed a significant deviant effect for rare deviant compared to standard stimuli. Second, in Experiment 2a, an ERP difference between standard and rare recombined stimuli was not significant either despite a much higher signal to noise ratio in comparison to Experiment 1b.

Our results provide evidence that crossmodal statistical relations are better implicitly learned in the developing than in the adult system. An enhanced sensitivity for low-level statistical patterns during development had been reported by other studies as well. For example, Janacsek et al. (2012) and Nemeth et al. (2013) demonstrated that children are superior in implicit statistical learning of sequences compared to adults but later loose this advantage and become more reliant on explicit learning. A similar developmental time course was found in a study of Jost et al. (2011), investigating the neurophysiological correlates of visual statistical learning in children and adults: children showed learning related ERP effects earlier during the acquisition phase indicating that they acquired the statistical structure quicker than the adult group. It is important to take into account that not all studies investigating statistical learning during development found enhanced learning performance in infants or children. For example, Saffran et al. (1996, 1999) reported similar abilities in eight-month-old infants and adults in the extraction of the underlying statistical structure of auditory sequences. Other studies observed better learning for older children and young adults than in younger age groups (Mayberry et al., 1995; Fletcher et al., 200; Kirkham et al., 2007). At first glance, these findings seem to be at odds with the present results. However, these inconsistent findings can be related to the complexity of the statistical patterns. Indeed, several studies have revealed that the ability to extract statistical patterns from sensory input during infancy improves from the simple tracking of event probabilities early in the development (from three months onwards, see Fantz et al., 1964) to the learning of more complex and higher-level statistical patterns at a later developmental stage (from twelve months onwards, see Gómez & Maye, 2005).

In addition to the enhanced sensitivity for crossmodal statistics in infants, our findings strongly suggest that learning mechanisms change from early development to adulthood. Adults did not learn crossmodal combinations implicitly as infants did, but succeeded when special crossmodal combinations were task relevant. Animal studies have suggested that during the sensitive phase, neural networks are set up in response to an exposure to the environment while during later development and in adulthood learning is context-specific and depends on task relevance (e.g. reward) and instructions (Keuroghlian & Knudsen, 2007). Currently, we can only speculate about the neural underpinnings of this age-dependent neuroplasticity. As noted by Dehaene-Lambertz & Spelke (2015) feedforward connectivity seems to be to a larger degree genetically determined than feedback connectivity and the latter seems to be mostly experience dependent. Changes in physical stimulus properties (in our study represented by rare deviant stimuli) can be detected to a larger extent based on feedforward connectivity and seems to work independent of task context both in infants and adults. This is in accordance with our results that infants as well as adults were able to differentiate the standard and rare deviant stimuli at an early processing stage. In contrast, the detection of rare recombined stimuli was associated with a longer latency ERP effect in both infants and adults. Indeed, multisensory binding has been found to rely on later processing stages in adults (Bruns & Röder, 2010a; Bonath et al., 2007). Emberson et al. (2011) provided evidence that crossmodal connectivity is at least partially in place at the age of six months. Here we speculate that this initial crossmodal connectivity might even be more extensive in the developing brain (see Johannsen & Röder, 2014) and thus might be the neural underpinning of the enhanced sensitivity to crossmodal statistics in development. We further assume in line with the “multisensory perceptual narrowing” idea (Lewkowicz & Ghazanfar, 2006) that experience narrows down the initial crossmodal connectivity and elaborates the connections which are useful for an individual (Johannsen & Röder, 2014; Lewkowicz, 2014). With an improved tuning of neural networks the learning mode changes towards a larger context dependency to guarantee the small adaptations necessary throughout life. Moreover, as some parts of the neural networks seem to stabilize, learning partially shifts to different neural sites. For example, while prism wearing during the sensitive phase changes the connectivity between the central (ICC) and external (ICX) inferior colliculus of the midbrain of barn owls, adaptation to prisms later in the critical period is mediated by a reorganization of the optical tectum to which the ICX projects (Knudsen, 2002). In the present study we found that the learning of crossmodal combinations in adults depends on task relevance (Experiment 2b). Thus, in accordance with Keuroghlian and Knudsen (Keuroghlian & Knudsen, 2007) and Bavelier et al. (2010), neuroplasticity in adults depends to a larger extent on task relevance and attention. Task relevance or attention constitute specific top-down influences on sensory representations and are thus mediated via feedback connections that were are well tuned and elaborated during development (Dehaene-Lambertz & Spelke, 2015).

Since we argue that the change in learning mode during development is related to functional specialization, the strong lateralization of both ERP effects in infants seems rather surprising. The differentiation of standard and rare recombined stimuli requires the detection of conditional probabilities. This ability has been postulated as a precursor of language learning. Indeed, it has been shown with structural imaging techniques that many hemispheric asymmetries, in particular those related to the language system (Friederici, 2009) exist at birth or shortly thereafter (see Dehaene-Lambertz & Spelke, 2015). Thus, we speculate that the strong left lateralized ERP difference between standard and rare recombined stimuli might reflect a recruitment of similar neural circuits that have been proposed to enable the detection of word boundaries (Saffran, 1996), non-adjacent transitional probabilities and possibly syntactical rules (Friederici, 2002; Friederici et al., 2006). Thus, this neural system might, partially independently of modality and domain, allow for detecting statistical relations (Kuhl, 2010; Aslin & Newport, 2014). The right lateralized ERP effect to rare deviant stimuli was not unique to the infant group, but was as well observed in the adults tested with the same passive design (Experiment 1b). Interestingly such a lateralization was neither found for Experiment 2a nor for Experiment 2b, in which the adult participants were actively engaged in a task. We speculate that rare deviants elicited a reflexive and exogenous attention shift to the rare sensory features. Such reflexive spatial attention orienting has often been associated with right parietal brain regions (Okada et al., 2008; Mort et al., 2003; Chica et al., 2011). In contrast, in Experiment 2a and 2b, participants had to allocate attention to a certain stimulus or stimulus combination and to avoid exogenous attention shifts.

In conclusion our study demonstrates that six-month old infants were able to quickly learn crossmodal statistics through a mere passive exposure, whereas adults learned the same crossmodal combinations only when they were task relevant. Thus, we provide first evidence for a higher sensitivity for crossmodal statistics in infants compared to adults, indicating age-dependent mechanisms for the learning of arbitrary crossmodal combinations. We speculate that initial passive association learning allows infants to quickly form first internal models of their sensory environment. In adulthood these internal models are adjusted if task relevant.

## Acknowledgments

This work was supported by the European Research Council (ERC-2009-AdG 249425 CriticalBrainChanges) and grant “Crossmodal Learning” of the City of Hamburg. We thank Rebecca Nixdorf for help with data acquisition and Erich Schröger for his comments and suggestions. We are particularly grateful to the parents and their children for taking part.

## Authors' contributions

All authors contributed to the design of the study; B.H., M.v.F., and S.R. collected the data; B.R., B.H. and S.R. analyzed the data; B.R. and S.R. wrote the manuscript; all authors checked and approved the final manuscript.

